# Investigating Mechanically Activated Currents from Trigeminal Neurons of Non-Human Primates

**DOI:** 10.1101/2024.10.06.616876

**Authors:** Karen A Lindquist, Jennifer Mecklenburg, Anahit H. Hovhannisyan, Shivani Ruparel, Armen N. Akopian

**Author notes:** **<u>Corresponding authors:</u>** Karen A. Lindquist University of Texas at Austin 100E 24th Street, Austin, TX 78712 Armen N. Akopian University of Texas Health Science Center @ San Antonio 7703 Floyd Curl Drive, San Antonio, TX 78229-3900.

## Abstract

**Introduction:** Pain sensation has predominantly mechanical modalities in many pain conditions. Mechanically activated (MA) ion channels on sensory neurons underly responsiveness to mechanical stimuli. The study aimed to address gaps in knowledge regarding MA current properties in higher order species such as non-human primates (NHP; common marmosets), and characterization of MA currents in trigeminal (TG) neuronal subtypes.

**Methods:** We employed patch clamp electrophysiology and immunohistochemistry (IHC) to associate MA current types to different marmoset TG neuronal groups. TG neurons were grouped according to presumed marker expression, action potential (AP) width, characteristic AP features, after-hyperpolarization parameters, presence/absence of AP trains and transient outward currents, and responses to mechanical stimuli.

**Results:** Marmoset TG were clustered into 5 C-fiber and 5 A-fiber neuronal groups. The C1 group likely represent non-peptidergic C-nociceptors, the C2-C4 groups resembles peptidergic C-nociceptors, while the C5 group could be either cold-nociceptors or C-low-threshold-mechanoreceptors (C-LTMR). Among C-fiber neurons only C4 were mechanically responsive. The A1 and A2 groups are likely A-nociceptors, while the A3-A5 groups probably denote different subtypes of A-low-threshold-mechanoreceptors (A-LTMRs). Among A-fiber neurons only A1 was mechanically unresponsive. IHC data was correlated with electrophysiology results and estimates that NHP TG has ∼25% peptidergic C-nociceptors, ∼20% non-peptidergic C-nociceptors, ∼30% A-nociceptors, ∼5% C-LTMR, and ∼20% A-LTMR.

**Conclusion:** Overall, marmoset TG neuronal subtypes and their associated MA currents have common and unique properties compared to previously reported data. Findings from this study could be the basis for investigation on MA current sensitizations and mechanical hypersensitivity during head and neck pain conditions.

## Introduction

Pain perception predominantly involves mechanical modalities in most acute and chronic pain conditions (Lolignier et al., 2015). The somatosensory system can distinguish between various non-painful mechanical sensations (Volkers et al., 2015). Neuronal circuits that discriminate between painful and non-painful mechanical sensations are anchored by specialized sensory neurons that respond to environmental mechanical stimuli. These neurons are housed in several sensory ganglia, including the trigeminal (TG) and dorsal root ganglia (DRG) (Hao and Delmas, 2011).

Somatosensory neurons are neurochemically and functionally diverse (Lumpkin and Caterina, 2007; Uceyler, 2016; Usoskin et al., 2015a). Beyond their classification by myelination status and conduction velocity (Basbaum and Braz, 2010), they can also be categorized based on their functional roles, such as nociceptors, mechanoreceptors, and proprioceptors. Traditionally, they are divided into unmyelinated C-fibers, thinly myelinated A-delta (Aδ), and myelinated A-beta (Aβ) fibers (Baumgartner, 2010), with further subdivisions into multiple subtypes (Basbaum et al., 2009; Djouhri et al., 1998a; Djouhri and Lawson, 2004). These distinct neuron types exhibit unique responses to mechanical stimuli. Mechanically activated (MA) ion channels underlie responses to proprioceptive stimuli, touch, and mechanical pain (Murthy et al., 2018; Tang et al., 2020). MA currents are sensitized in pain conditions and contribute directly to mechanical hypersensitivity (Bhave and Gereau, 2004; Dubin et al., 2012; Hucho and Levine, 2007; Schaefer et al., 2023). Understanding the relationship between MA currents and specific sensory neuron subtypes in various pain conditions and stages of chronic pain is critical for a more comprehensive understanding of pain mechanisms.

MA currents can be elicited and recorded using various techniques, one of approaches to activate MA involves a piezo-actuator device, originally designed for patch-clamp recording of MA in DRG neurons (McCarter et al., 1999). Distinct MA currents have since been recorded in mouse DRG neurons (Coste et al., 2010; Drew et al., 2002). Using this approach, numerous studies have demonstrated how different agents modulate MA currents (Di Castro et al., 2006; Dubin et al., 2012; Narayanan et al., 2018; Romero et al., 2023; Romero et al., 2020; Zhang et al., 2019). However, significant gaps in our understanding remain. First, most data have been gathered from DRG neurons, while TG neuron MA currents remain largely unexplored. To date, TG neuron MA currents have mainly been studied in birds (Schneider et al., 2019; Schneider et al., 2017; Ziolkowski et al., 2023) and in mouse TG neurons innervating the cornea (Fernandez-Trillo et al., 2020). Second, MA currents in human sensory neurons have been examined using iPSC-derived neurons rather than naïve human sensory neurons (Romero et al., 2020; Schrenk-Siemens et al., 2015), which may not fully represent the properties of mature sensory neurons. Third, there is limited data on the relationship between MA currents and specific sensory neuron types. Notable exceptions include studies examining MA currents in skin and muscle-innervating DRG neurons (Weyer et al., 2015) and TG neurons innervating the cornea (Fernandez-Trillo et al., 2020).

We investigated the properties of MA currents in TG neurons, using non-human primates (NHPs), specifically common marmosets, to provide data that are evolutionarily closer to humans. Additionally, we associated MA currents with specific sensory neuron types. Given the impracticality of using reporter animals in NHPs, we classified sensory neuron types based on electrophysiological parameters, including action potential (AP), after-hyperpolarization (AHP) and AP train (Lindquist et al., 2021; Patil et al., 2018; Petruska et al., 2000a). The data generated here aim to fill these knowledge gaps, enhance the translatability of findings, and, when combined with transcriptomic data, offer insights into the signaling pathways that modulate MA currents and, ultimately, the peripheral mechanisms underlying mechanical orofacial pain.

## Methods

### Ethical Approval

This study adheres to the ARRIVE 2.0 guidelines (Percie du Sert et al., 2020). All animal care and experimental procedures complied with the United States Public Health Service Policy on Humane Care and Use of Laboratory Animals, the Guide for the Care and Use of Laboratory Animals, and the American Society of Primatologists’ principles for the ethical treatment of NHP. We followed guidelines from the National Institutes of Health (NIH) and the Society for Neuroscience (SfN) to minimize the number of animals used and their suffering. All procedures were approved by the Institutional Animal Care and Use Committee (IACUC) of the University of Texas Health Science Center at San Antonio (UTHSCSA) and the Texas Biomedical Research Institute (TBRI). The IACUC protocol titles are “Plasticity of lymphotoxin-beta signaling and orofacial pain in non-human primates” (UTHSCSA: 20200021AR; TBRI: 1821 CJ 0).

### Animals subjects, tissue collection and transfer

We collected tissues from five adult male common marmosets (Callithrix jacchus), aged 3-5 years. Tissue samples were obtained at necropsy from the UTHSCSA and TBRI “Tissue Share” program, where animals were euthanized under IACUC-approved endpoints. None of the animals had head or neck injuries or systemic infections. Following euthanasia by veterinary staff at either UTHSCSA or TBRI, trigeminal ganglia (TG) were collected within 2 hours of death. Tissues were preserved for patch-clamp electrophysiology in Hank’s solution on ice or fixed in 4% paraformaldehyde (PFA) for immunohistochemistry (IHC) as previously described (Ibrahim et al., 2023). Samples were transported by car (15-minute drive) from TBRI to UTHSCSA.

### Primary Trigeminal Ganglion Neuronal Culture

TG cultures were initiated immediately upon sample arrival. Tissues were incubated for 45 minutes in Hank’s solution containing 30 µL of 50 mg/mL Collagenase I (Worthington, Code #CLS-1, 230U/mg) and 7.5 µL of 50 mg/mL Dispase (Roche, Catalog #4942078001). Tissues were washed twice with DMEM (with 4% fetal bovine serum, 2 mM L-glutamine, 100 U/mL penicillin, 100 µg/mL streptomycin) and centrifuged at 1000 RPM for 75 seconds. Cells were mechanically dissociated and plated on 12 mm German glass coverslips coated with laminin and poly-D-lysine (Corning, Catalog #08-774-385). No growth factors were added to the media to prevent artificially enhancing neuronal responses. This is a key point because growth factors like NGF are known to sensitize sensory neurons leading to artificially enhanced MA currents and unmask silent nociceptors (Prato et al., 2017; Price et al., 2005; Ritter and Mendell, 1992; Schaefer et al., 2018; Shrivastava et al., 2021). Electrophysiological experiments were conducted 6-24 hours after plating.

### Whole-cell patch clamp electrophysiology

Recordings were performed at 22-24°C using an Axopatch 200B amplifier and analyzed with Axon pClamp11.2 software (Molecular Devices). Data were filtered at 0.5-5 kHz and sampled at 2-20 kHz depending on current kinetics. Fire-polished glass electrodes (3–7 MΩ resistance) were used. Access resistance (Rs) was compensated (40-80%) when appropriate. Data were discarded if Rs changed >20% during recording, leak currents were >100 pA, or input resistance was <100 MΩ. Currents were considered positive when amplitudes were at least 5-fold larger than noise levels. The standard external solution (SES) contained 140 mM NaCl, 5 mM KCl, 2 mM CaCl_2_, 1 mM MgCl_2_, 10 mM D-glucose, and 10 mM HEPES (pH 7.4). The standard pipette solution (SIS) contained 140 mM KCl, 1 mM MgCl_2_, 1 mM CaCl_2_, 10 mM EGTA, 10 mM D-glucose, and 10 mM HEPES (pH 7.3) along with 2.5 mM ATP and 0.2 mM GTP.

### Electrophysiology protocols, MA current recording, and data analysis

After patching a selected neuron, recordings were made using a sequence of protocols: 1) In the current clamp configuration, a single AP response was generated with a 1, 2, 3, 4 or 5nA (separate sweeps) 0.5ms current pulse; 2) Following this, AP trains were elicited by applying step currents of 50–550pA with 100-pA increments for 1s long sweeps or 200–2000pA with 300pA increments for the few >80-100pF neurons that did not fire a single AP; 3) Electronics were then switched to voltage-clamp configuration (V_hold_ at -60mV) and MA currents were recorded (elaborated below); and 4) the final protocol in voltage clamp configuration for outward I_A_ K+ currents was a step down from Vh to -70mV, kept for 500ms, and then 200ms depolarizing command steps (20 mV) applied from -40mV to a final potential of +20mV (Lindquist et al., 2021; Patil et al., 2018; Petruska et al., 2000b). The following variables were measured: cell capacitance (C_m_ in pF); resting membrane potential (RMP in mV), AP width at a RMP level (dB in ms), characteristic features of AP shapes, AHP peak (in mV), AHP_80_ duration (in ms), threshold of activation for MA currents (in actuating pipette displacement μm), their peak current densities, τ (time constant in msec) decay for MA current inactivation kinetics using the equation I =ΔI*exp^(-t/τ_inact_) (Petruska et al., 2000a), and from the outward current protocol-4, the trace evoked by +20 mV was fit with a standard (i.e single) exponential function (A_1_ exp[-(t-k)/τ] + C). Fitting and decay tau (τ; ms) calculation was performed using pCLAMP11.2 software (Molecular Devices, Sunnyvale, CA). Additionally, presence or absence of I_A_-current peaks in protocol-4 at steps to 0 and +20 mV was an important variable for grouping of some neurons. Thus, presence of clear transient potassium current (i.e. I_A_) classified DRG neuron as a A-fiber (Patil et al., 2018; Petruska et al., 2000b; Usoskin et al., 2015a). C_m_ can be recalculated into cell diameter using d=5*√(Cm/4π); where Cm is capacitance in pF, d is diameter in μm, if neurons are considered spherical.

Deformation of the plasma membrane produced by mechano-clamp is analogous to mechanical indentation *in vivo* (Hao and Delmas, 2011). In a voltage-clamp (V_hold_=-60mV) configuration, MA currents were induced by mechanical stimulation which was directly applied by a fire-polished blunt borosilicate glass pipette (BF100-50-10, Sutter Instruments) with a tip diameter of 1-3µm, held at a 45° angle from the coverslip. This pipette was driven by a piezoelectric actuator (P-841.6, Physik Instrumente) (Hao and Delmas, 2011; McCarter et al., 1999). Probe movement was controlled by a Digital Piezo Controller (E-709, Physik Instrumente) which in turn was under control of Axopatch 200B amplifier and Clampex 11.2 software (Molecular Devices, Sunnyvale, CA). The tip was advanced in increasing 1.5µm displacements, each held for 300ms, with 10 sec between steps to avoid sensitization or desensitization of currents during 10 consecutive sweeps (Weyer et al., 2015). If mechanically responsive, current amplitudes incrementally increased as each mechanical poke deepened (Rugiero et al., 2010).

### Immunohistochemistry

Immunostaining of marmoset TGs was performed as described previously (Hovhannisyan et al., 2023; Tram et al., 2023). TG tissues were fixed in 4% PFA for at least 16 hours, cryoprotected with 10% and 30% sucrose in PBS for at least 24 hours, embedded in Neg-50 (Richard Allan Scientific, Kalamazoo, MI), and cryosectioned at 20 μm thickness. Sections were blocked with 4% normal donkey serum (Sigma, St. Louis, MO), 2% bovine gamma-globulin (Sigma-Aldrich, St. Louis, MO) and 0.3% Triton X-100 (Fisher Scientific) in 0.1M PBS for 90 min at RT, and subsequently incubated with primary antibodies for 24-36 hours. Sections were then washed with PBS from unbound primary antibodies, blocked, and incubated for 90 min at RT with appropriate fluorophore conjugated secondary antibodies (Jackson Immuno-Research, West Grove, PA, USA). Finally, tissue sections were washed for 3 x 5 minutes with 0.1M PBS and 2 x 5 minutes in diH_2_O, air-dried, and covered with Vectashield Antifade Mounting Medium (Vectorlabs, Burlingame, CA, USA). The following previously characterized primary antibodies were used: anti-CGRP guinea pig polyclonal (Synaptic Systems; Goettingen, Germany; catalog 414 004; 1:200) (Samms et al., 2022); anti-TRPV1 rabbit polyclonal (Novus Biologicals; Centennial, CO; catalog NBP1-71774SS; 1:200) (Moutafidi et al., 2021); anti-mrgprD rabbit polyclonal (Alamone Lab; catalog ASR-031; 1:200) (de Carvalho Santuchi et al., 2019; Lindquist et al., 2021); anti-tyrosine hydroxylase (TH) chicken polyclonal (Neuromics; Bloomington, MN; catalog CH23006; 1:300) (Ferreira-Pinto et al., 2021); anti-trkC rabbit polyclonal (Aviva Systems Biology, San Diego, CA; catalog ARP51318_P050; 1:200); anti-trkB goat polyclonal (R&D systems; AF1494; 1:200) (Arcourt et al., 2017); anti-parvalbumin rabbit polyclonal (Novus Biologicals; catalogue NB120-11427SS; 1:200) (Lewis et al., 2021); and anti-calbindin D28k rabbit polyclonal (Synaptic Systems; catalogue 214 011; 1:200) (Navarro-Gonzalez et al., 2021).

Z-stack images were captured with a Keyence BZ-X810 all-in-one microscope (Itasca, IL, USA) under the ‘sectioning’ mode using a 2×, 10× or 20× objective. Control IHC was performed on tissue sections processed as described but either lacking primary antibodies or lacking primary and secondary antibodies. Settings were determined in such way that no-primary antibodies and both no-primary and no-secondary antibody controls did not show any positive signal. Then, images were taken using these fixed acquisition parameters across all groups. For cell counting, Z-stack IHC images with x10 or x20 objectives were obtained from 3-5 independent tissue sections from 2-3 primates/isolations. Counted marker-positive neurons were presented as percentages of the total neuron numbers. All TG could have been labeled by pan sensory neuronal marker (NeuN) (Mecklenburg et al., 2023; Wu et al., 2018). However, NeuN did not clearly labeled at least 25% NHP TG neurons, and use polyclonal rabbit-made NeuN could compromised usage of many available antibodies. Accordingly, the total neuronal numbers were estimated by taking section pictures with high gain and highlighting auto-fluorescence in all TG neurons. Neurons were counted using Image J software (Tram et al., 2023). Mean values from counting from 3-5 sections generated from a NPH TG represented data for a biological replicate. Thus, n=3 means represent 3 NHP TG as the biological replicates.

### Statistical Analysis

GraphPad Prism 10 (GraphPad) was used for statistical analyses. Data in the figures are mean ± SEM, with *n* referring to number of sectioned TG for IHC and the numbers of analyzed recorded cells for electrophysiology. Differences between expression percentages of markers in TG and electrophysiologically characterized groups were assessed by unpaired *t*-test or one-way ANOVA with Tukey’s post hoc tests, each column was compared with all other columns. A difference is accepted as statistically significant when p < 0.05. Interaction F ratios, and the associated p values are reported.

## Results

### Clustering TG neurons according to electrical and MA current properties

Sensory neuron classification can be achieved through multiple approaches, including transcriptomic profiling (Usoskin et al., 2015b; Yang et al., 2022), immunohistochemistry, reporter animal data, *ex vivo* or *in vivo* extracellular recordings, and electrophysiological properties derived from patch-clamp recordings (Boada, 2013; Lindquist et al., 2021; Patil et al., 2018; Petruska et al., 2000a), or a combination of these methods (Zheng et al., 2019). In rodent models, sensory neuron size is a dependable indicator for differentiating between DRG C-and A-fiber neurons (Lindquist et al., 2021; Patil et al., 2018). However, in NHP TG neurons, this distinction based on size (capacitance) was less clear (Table 1). Many NHP TG neurons exceeded a capacitance of 100 pF, which is the upper limit for the Axopatch 200B amplifier.

**Table 1:** Electrophysiological properties of common marmoset TG neuronal groups.

| Group | N | Cap (pF) | RMP (mV) | dB (ms) | AP Shape | AHP<sub>80</sub> (ms) | AHP peak (mV) | AP Thresh (pA) | AP Train (Hz) | K<sup>+</sup> $\tau$ (ms) | MA (Y/N) |
| --- | --- | --- | --- | --- | --- | --- | --- | --- | --- | --- | --- |
| C1 | 7 | <sup>1</sup>69.4 ± 7.7 | -54.1 ± 2.8 | 11.1 ± 0.9 | Bow | 23.9 ± 12 | -9 ± 2.2 | N/A | N/A | N/A | N |
| C2 | 20 | <sup>1</sup>66.3 ± 5.3 | -49 ± 2.1 | 10 ± 0.7 | Hump | 30.4 ± 6.8 | -9.4 ± 0.8 | N/A | <sup>3</sup>N/A | <sup>5</sup>9.1 ± 0 | N |
| C3 | 29 | 62.6 ± 3.7 | -56.4 ± 1.7 | 8.1 ± 0.3 | Hump | 48.1 ± 7.5 | -11.4 ± 0.7 | 125.9 ± 12.8 | 8.4 ± 1.8 | 13.1 ± 3.3 | N |
| C4 | 12 | <sup>1</sup>59.1 ± 5.6 | -63.4 ± 3.5 | 10.7 ± 1.1 | Hump | 32.7 ± 9.1 | -7.5 ± 1 | 161.1 ± 20 | 4.6 ± 0.8 | <sup>5</sup>14 ± 0 | Y |
| C5 | 15 | <sup>1</sup>61.2 ± 5.9 | -38.3 ± 2.4 | 7.6 ± 0.9 | <sup>2</sup>Straight | 8 ± 3.1 | -5.5 ± 0.8 | N/A | <sup>3</sup>N/A | <sup>5</sup>5.4 ± 0 | N |
| A1 | 32 | <sup>1</sup>89.7 ± 3.2 | -62.8 ± 1.5 | 5.5 ± 0.3 | Deflection | 48.7 ± 7.1 | -10.3 ± 0.9 | 173 ± 28.1 | 8.4 ± 2.7 | 10.5 ± 1.3 | N |
| A2 | 21 | <sup>1</sup>96.3 ± 2.6 | -63.6 ± 1.5 | 4.6 ± 0.4 | Deflection | 55 ± 11.3 | -9.6 ± 1.2 | 175 ± 25 | 5.5 ± 1.7 | 10.2 ± 0.9 | Y |
| A3 | 9 | <sup>1</sup>79.6 ± 7.1 | -68.3 ± 1.9 | 2.6 ± 0.1 | Straight | 1.7 ± 0.9 | -4 ± 0.4 | N/A | <sup>3</sup>N/A | 4.3 ± 0.4 | Y |
| A4 | 15 | <sup>1</sup>78.7 ± 5.5 | -64.4 ± 1 | 2.6 ± 0.1 | Straight | 41.9 ± 10.6 | -7.8 ± 0.9 | N/A | <sup>3</sup>N/A | 5.5 ± 0.4 | Y |
| A5 | 25 | <sup>1</sup>71.6 ± 4.5 | -59 ± 2.3 | 2.3 ± 0.1 | Straight | 11.8 ± 1.9 | -9.4 ± 0.9 | N/A | <sup>4</sup>N/A | 4.8 ± 0.3 | <sup>6</sup>Y |

Values are means ± SEM. AP parameters, $I_A \tau$ , and whether the neuronal subclass is mechanically responsive is presented. AP threshold was determined for neurons that fired multiple APs. <sup>1</sup>Inaccurate average capacitance because neurons greater than 100pF could not be measured. C1 had 1, C2 had 3, C4 had 1, C5 had 1, A1 had 12, A2 had 18, A3 had 2, A4 had 5, and A5 had 4 neurons larger than 100pF. <sup>2</sup>Unique slightly inward curving shape. <sup>3</sup>One neuron from the group fired 2 APs. <sup>4</sup>Two neurons fired small trains of APs. <sup>5</sup>Only 1 $I_A$ current was recorded from each subgroup. <sup>6</sup>Several cells broke down mid-recording and have no MA recordings and another 2 did not respond to mechanical stimulation.

In our study, we classified NHP TG neurons based on electrophysiological parameters obtained via patch-clamp recordings for MA currents. Extensive literature supports the use of such recordings to reliably distinguish between sensory neurons containing C-and A-fibers (Lindquist et al., 2021; Petruska et al., 2000a; Zheng et al., 2019). Thus, for sensory neuronal classification, we utilized action potential (AP) shape, as previously established in our studies on mice (Patil et al., 2018). Neurons with broad APs (duration at base (dB) > 5-6 ms) and a characteristic “hump” on the downward slope of the AP (*Fig. 1A*) were classified as peptidergic C-high threshold mechanoreceptors (C-HTMRs; aka nociceptors). The presence of the “bow” suggests that these APs likely originate from non-peptidergic C-fiber nociceptors (Patil et al., 2018). Neurons with broader APs but without the hump could be categorized as cold-nociceptors or C-low threshold mechanoreceptors (C-LTMRs) (Patil et al., 2018). TG neurons with an AP duration of 2-5 ms and a distinct “deflection” on the downward slope were classified as A-HTMRs (Patil et al., 2018). Finally, neurons with fast APs (duration 1-3 ms) and a straight downward slope were classified as A-LTMRs. These main classes were further subdivided based on afterhyperpolarization (AHP) peak, AHP recovery time (AHP_80_; *Fig. 1A*), and the ability to generate AP trains (*Fig. 1B*). Neurons were also classified as mechanically sensitive and insensitive based on currents obtained upon piezo application (*Fig 1C*). Additionally, parameters related to outward currents, which are associated with transient A-type (I_A_) potassium (K⁺) currents (*Fig 1D*), were not used as a classification criterion due to technical limitations related to cells deteriorated before the final mechanical poke, making it impossible to collect outward current data from all recorded neurons.

**Figure 1.**
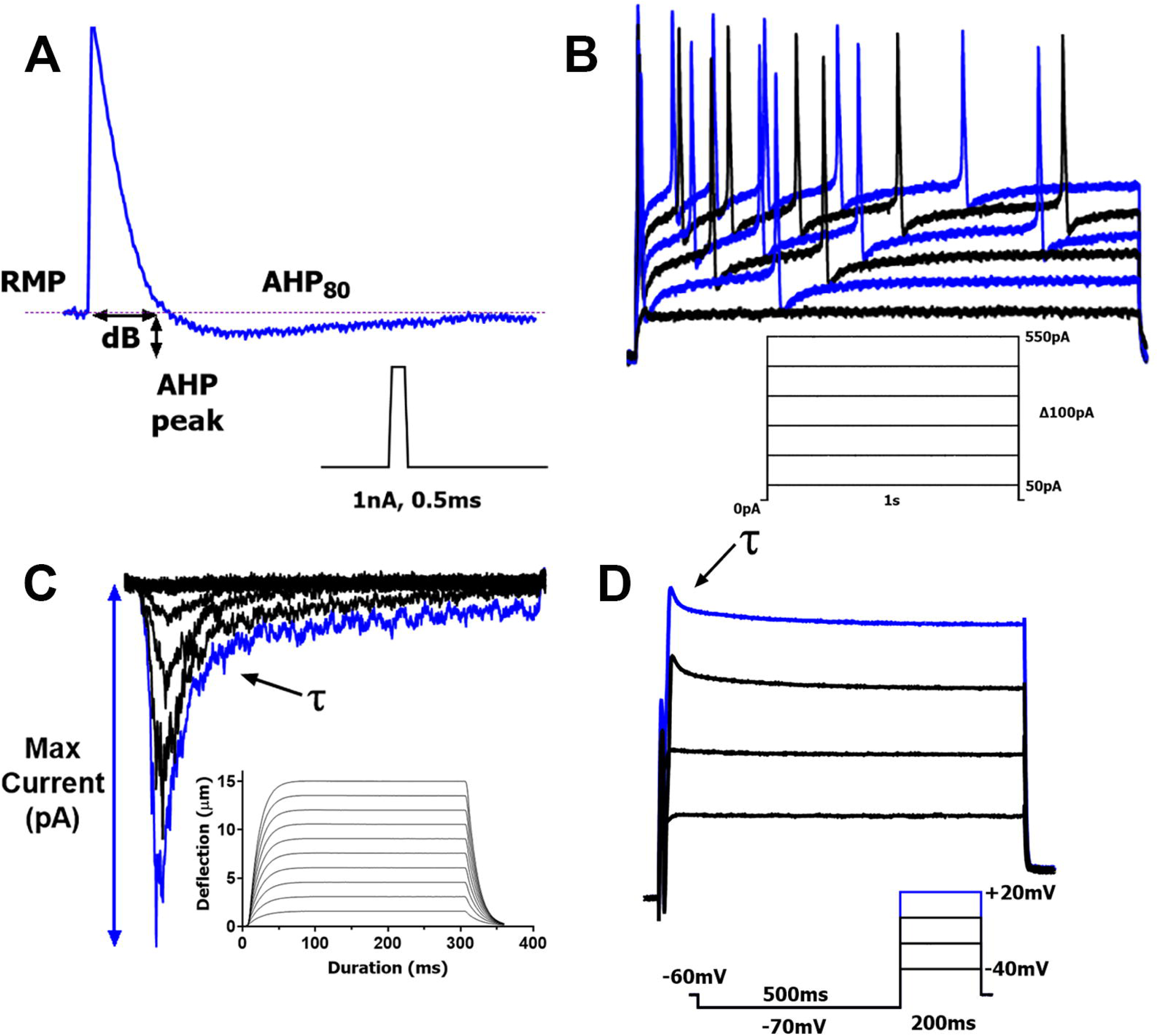
Electrophysiology protocols. **(A)** A single action potential (AP) is elicited by a brief current injection of 1-5 nA for 0.5ms. Stimulus waveform (1-5 nA, 0.5 msec) is indicated below, a trace generates a single AP in a NHP TG neuron. We analyzed indicated AP and afterhyperpolarization (AHP) parameters, including resting membrane potential (RMP); duration at base (dB), magnitude of AHP peak; and the time required for the AHP (measured in ms) to decay by 80% (AHP_80_). **(B)** AP trains are triggered by applying steps of increasing current injections, 50-550pA with 100pA increments. A schematic of the protocol used, and a sample recording is shown. **(C)** MA currents are activated by a Piezo actuator controlled by a piezo-electric device. Graphical representation of actuator movement is shown. Each poke extends by 1.5µm increments and held for 300ms before returning to the starting position for 10s of relaxation. A total of 10 progressively deeper pokes are administered, each increasing by an additional 15µm with the final poke going in 15µm deep. Sample recording from a NHP TG neuron is shown, τ is indicated. **(D)** Transient outward A-type current (I_A_) was generated by the indicated waveforms found below traces. I_A_ current is elicited by increasing positive voltage steps from -40mV to +20mV each held for 200ms in 20mV intervals after 500ms of a hyperpolarizing step down to -70mV from V_hold_. The decay constant τ was derived from a standard single exponential fitting between points indicated by arrows for the final outward current trace (+20 mV). Characteristic A-current peak is indicated by an arrow on the final trace generated by stepping to +20 mV.

More than 200 NHP TG neurons were recorded, and 185 common marmoset TG neurons that passed quality control were included in the analysis. Using the electrophysiological classification techniques described, we identified 5 distinct groups of C-fiber and 5 groups of A-fiber-containing TG neurons. The characteristics of these neuronal groups are detailed in *Table 1*. Of the recorded neurons, 72 out of 185 responded to mechanical stimulation, which is consistent with previously reported proportions for mouse TG neurons innervating the cornea (Fernandez-Trillo et al., 2020). Of the 10 identified NHP TG neuron groups, a C-fiber subgroup (C4) and 4 A-fiber subtypes (A2-A5) responded to piezo-stimulation, displaying mechanically activated (MA) currents with varying magnitudes and activation thresholds (*Figs 2A, 2B; Table 2*). We did not classify MA currents based on their kinetics (rapid, intermediate, or slow) (Coste et al., 2010; Hao and Delmas, 2010; McCarter et al., 1999), as no significant differences were observed in the decay kinetics between the MA currents of the C4 and A2-A5 groups (1-way ANOVA; F (4, 64) = 1.216; P=0.3126; *Fig 2C; n=8-21; Table 2*). Additionally, MA kinetics can be an unreliable classification parameter, as it is highly dependent on the mechanical stimulation setup, including the piezo-actuator speed and approach angle .

**Figure 2.**
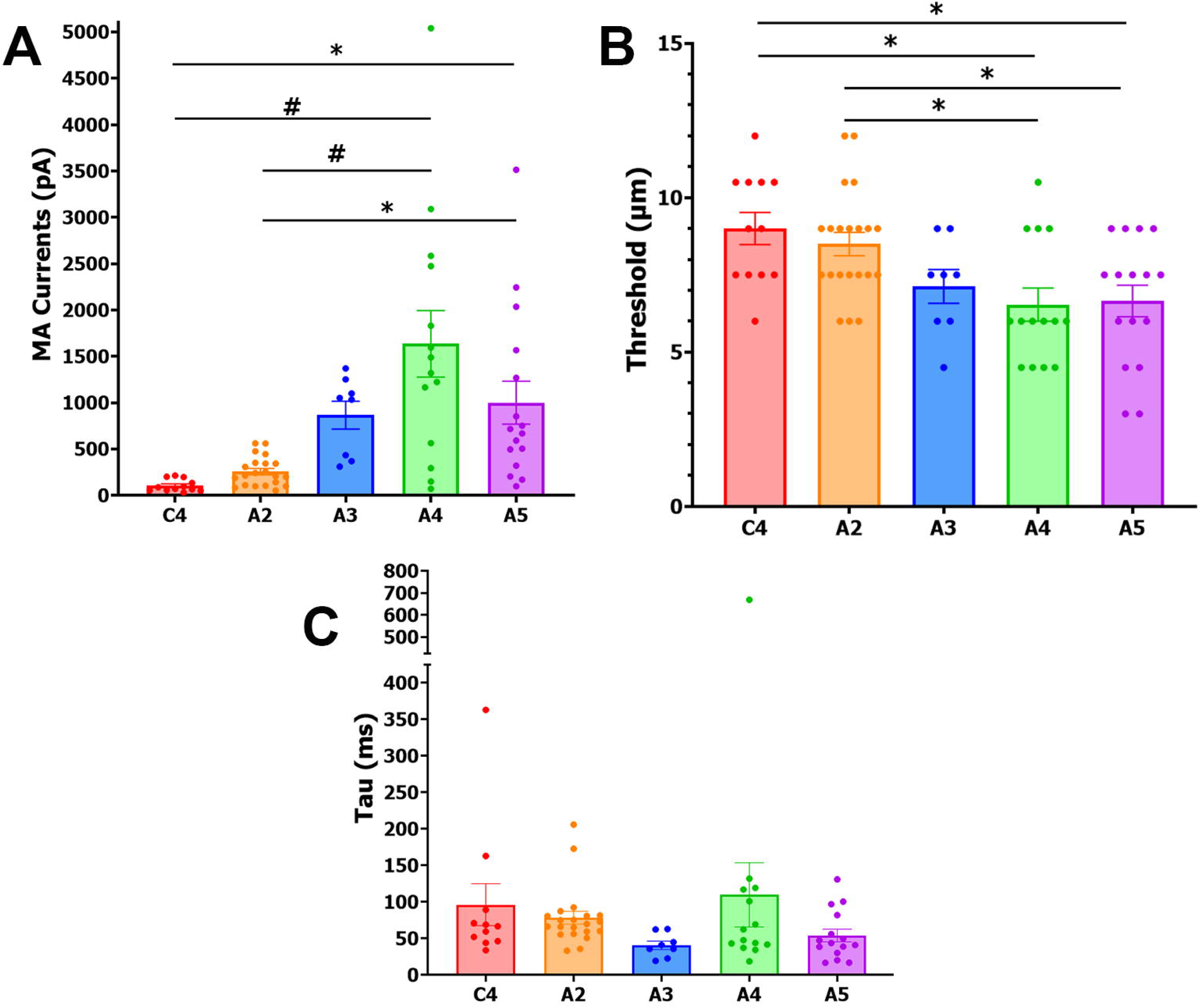
MA currents from NHP TG neurons. Five NPH neuronal subgroups (labeled under X-axis) responded to mechanical stimulation. These neuronal groups showed different characteristics. **(A)** Max MA current amplitudes (pA) from theses 5 MA groups of NHP TG neurons. **(B)** Activation threshold in actuator distance (μm) traveled for these NPH TG neuronal subgroups. **(C)** Decay kinetics (ms) of MA currents for these NPH TG neuronal subgroups. Data was analyzed by one-way ANOVA each column compared to others followed by Bonferroni’s post hoc tests; * - p<0.05; # - p<0.0001.

**Table 2:**
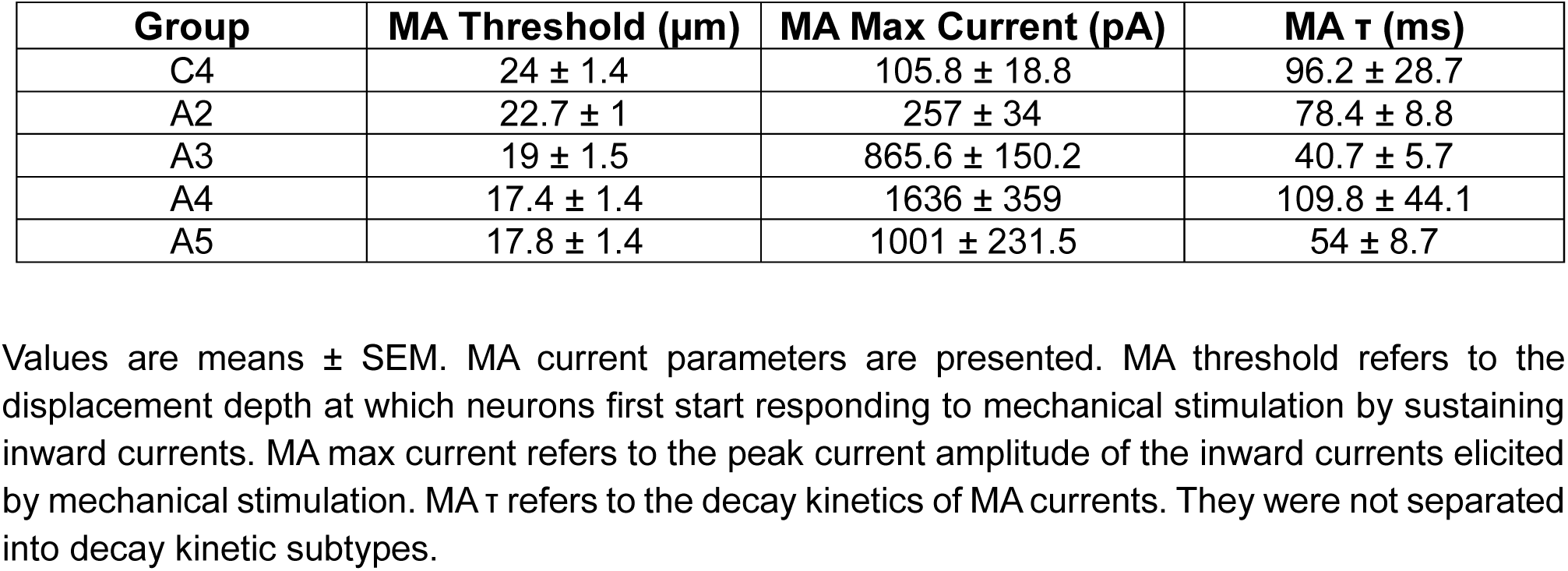
Properties of MA currents

### C-fiber TG neurons in common marmosets

This section outlines the distinct properties of each C-fiber-containing NHP TG neuron subgroup. Resting membrane potential (RMP) served as the primary quality control, with a cutoff range above -40 mV, as established in previous literature (Djouhri et al., 1998b; Fang et al., 2005). Neurons that met these criteria were included in the analysis.

<u>C1 Neurons</u>: The C1 group was the only subset of TG neurons exhibiting “bow”-shaped action potentials (APs) on the downward phase, with the largest AP dB among all C-fibers (*Fig 3A*). These neurons did not fire trains of APs nor respond to mechanical stimulation. C1 neurons had the largest soma size among C-fiber-containing neurons, as indicated by capacitance measurements (*Table 1*). This group was the least abundant of all recorded TG neurons. In mouse models, DRG and TG neurons with these properties—particularly the “bow”-shaped AP—are often associated with non-peptidergic, MrgprD+ C-fiber neurons (Lindquist et al., 2021; Patil et al., 2018).

**Figure 3.**
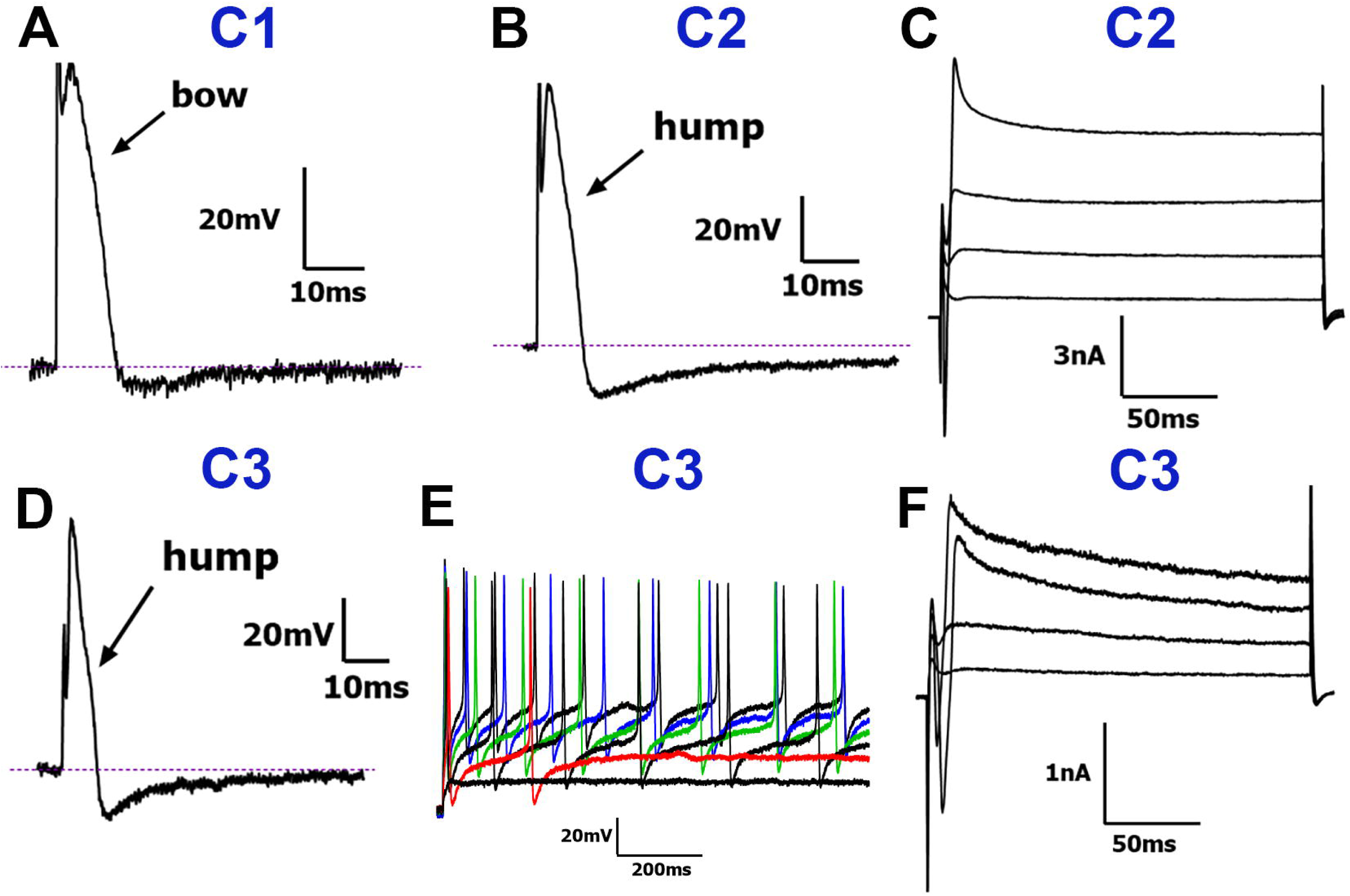
Traces representing current signatures from C1-C3 NPH TG neuronal groups. **(A)** Representative AP trace belonging to the C1 group. **(B)** Representative AP belonging to the C2 group. **(C)** Representative I_A_ belonging to the C2 group. **(D)** Representative AP belonging to the C3 group. **(E)** Representative AP train belonging to the C3 group. **(F)** Representative I_A_ belonging to the C3 group. A characteristic “hump” or “bow” on the downward portion of the AP is indicated with a black arrow in *panels A, B and D*. Neuronal groups are specified above traces. Scale bars are presented for each panel.

<u>C2 Neurons</u>: The C2 group exhibited long-duration APs with a characteristic “hump” on the falling phase (*Fig 3B*). Like C1 neurons, they did not fire AP trains or respond to mechanical stimulation. Only one neuron from this group had a complete set of recordings, including outward I_A_ currents (*Fig 3C*).

<u>C3 Neurons</u>: The C3 group, also characterized by a “hump” on the falling phase of the AP (*Fig 3D*), was the most abundant type of C-fiber. These neurons consistently fired AP trains and had the lowest AP firing threshold (*Fig 3E, Table 1*). Among C-fibers, I_A_ currents were reliably recorded only from this group (*Fig 3F*).

<u>C4 Neurons</u>: C4 neurons, which also had a “hump” on the falling phase of the AP (*Fig 4A*), were notable for their mixed firing behavior: most fired AP trains, but 3 out of 12 neurons fired only a single AP (*Fig 4B*). Only one I_A_ current recording was obtained from this group (*Fig 6C*). Importantly, C4 was the only C-fiber subgroup that exhibited MA currents (*Fig 6D*). However, compared to the other MA current-exhibiting subgroups, C4 neurons had the highest activation threshold and produced the smallest MA current amplitudes, though these values were not significantly different from those of A2 neurons (1-way ANOVA; F (4, 66) = 9.596; n=12 and 21; *Table 2, Fig 2A*). The C2-C4 groups display features consistent with C-fiber peptidergic neurons (Lindquist et al., 2021; Patil et al., 2018).

**Figure 4.**
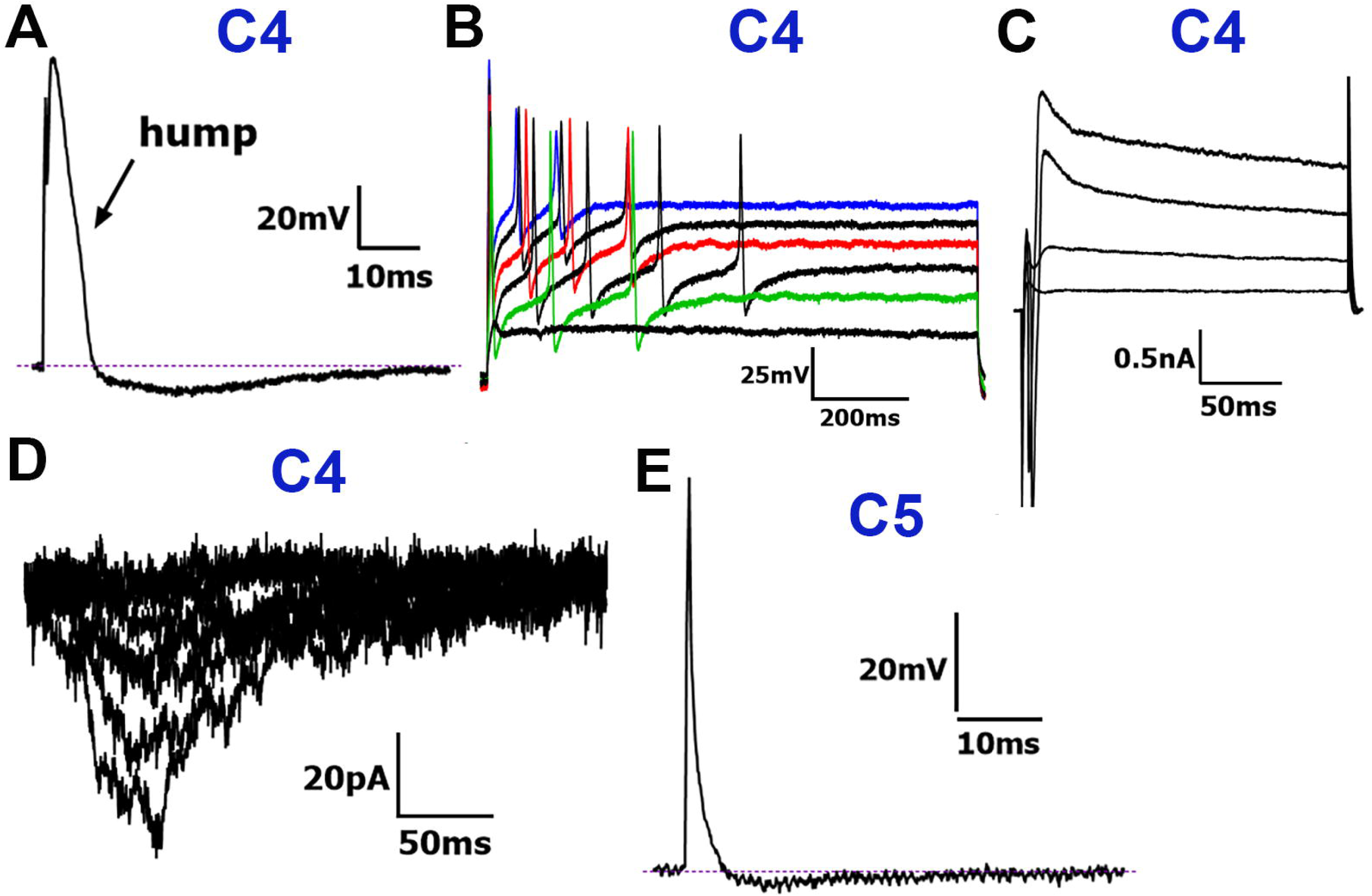
Traces representing current signatures from C4 and C5 NPH TG neuronal groups. **(A)** Representative AP trace belonging to the C4 group **(B)** Representative AP train belonging to the C4 group. (**C**) Representative I_A_ belonging to the C4 group. **(D)** Representative MA currents belonging to the C4 group. **(E**) Representative AP trace belonging to the C5 group. Neuronal groups are specified above traces. Scale bars are presented for each panel.

**Figure 5.**
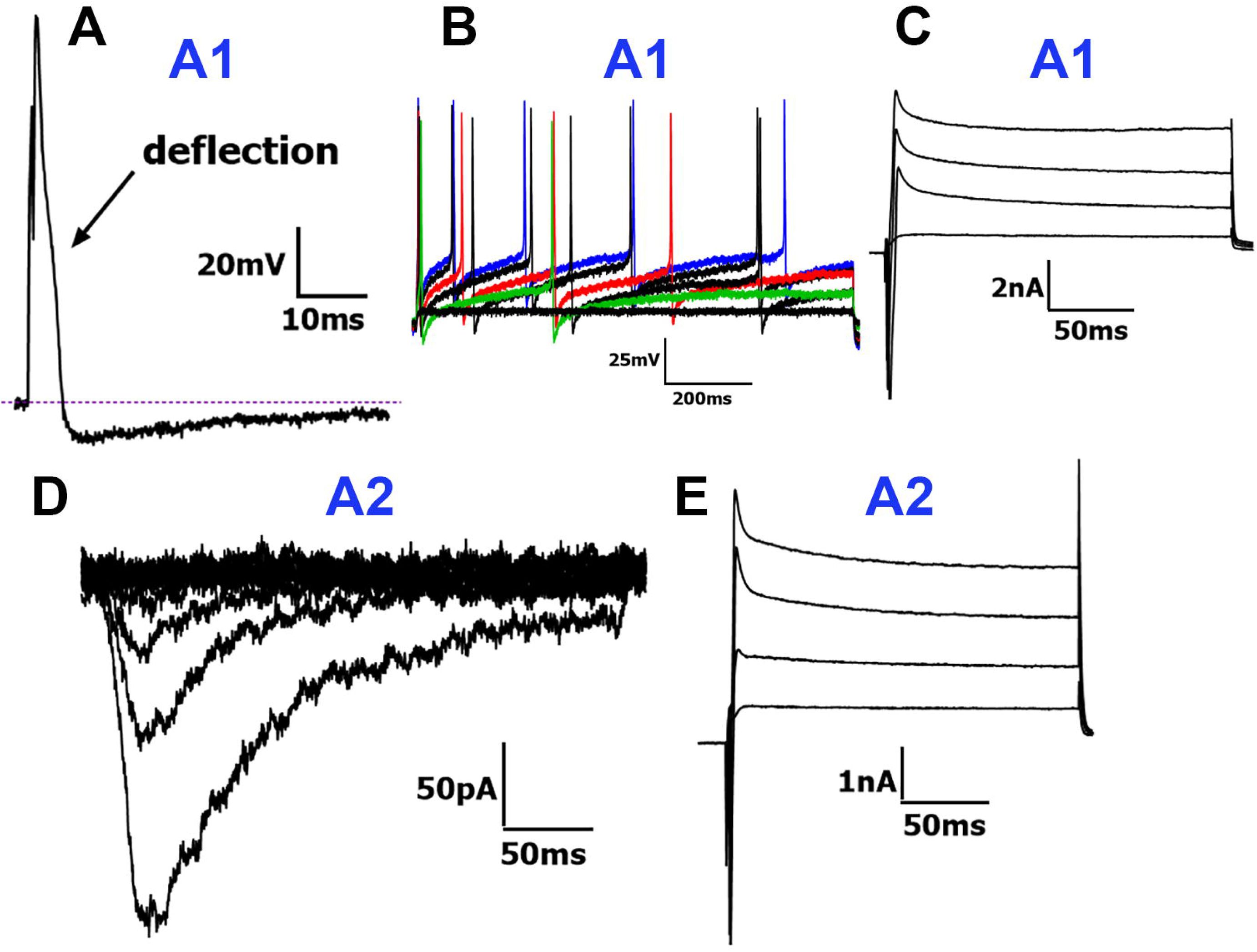
Traces representing current signatures from A1 and A2 NPH TG neuronal groups. **(A)** Representative AP trace belonging to the A1 group. Characteristic “deflection” on the downward part of the AP is indicated. **(B)** Representative AP train belonging to the A1 group. **(C)** Representative I_A_ belonging to the A1 group . **(D)** Representative MA currents belonging to the A2 group. **(E)** Representative I_A_ belonging to the A2 group. Neuronal groups are specified above traces. Scale bars are presented for each panel.

**Figure 6.**
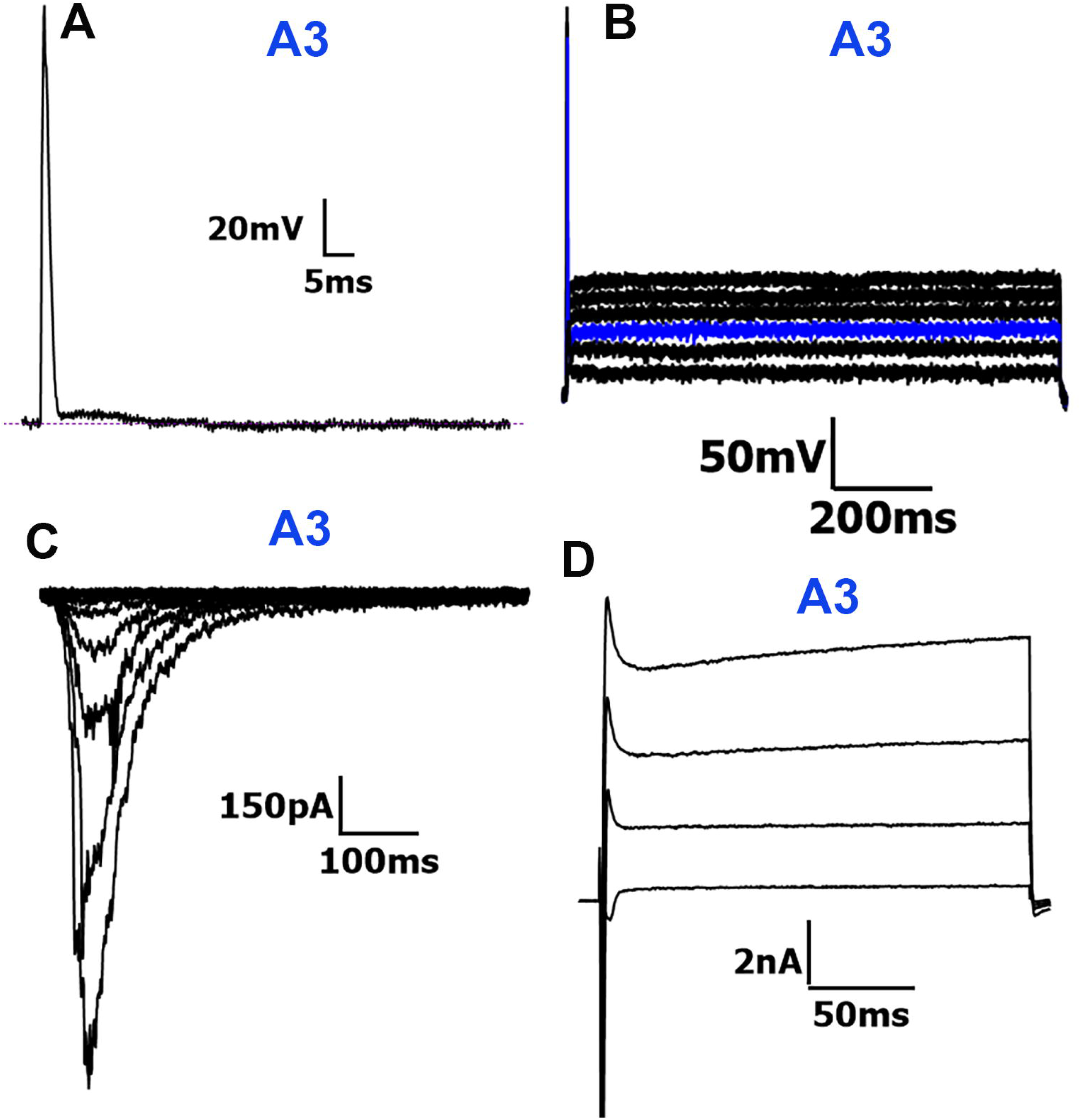
Traces representing current signatures from the A3 NPH TG neuronal group. **(A)** Representative AP trace belonging to the A3 group . “Afterdepolarization” in the AHP leading to the absence of a true AHP peak (see Fig 3A) is observed. **(B)** Representative AP train belonging to the A3 group . **(C)** Representative MA currents belonging to the A3 group. **(D)** Representative I_A_ belonging to the A3 group. Neuronal groups are specified above traces. Scale bars are presented for each panel.

<u>C5 Neurons</u>: The C5 group was the only C-fiber subgroup with long-duration APs that lacked inflections on the falling phase (*Fig 4E*). These neurons had the smallest AP dB among C-fibers, though still longer than that of A-fibers (*Table 1*). They also exhibited the fastest AHP_80_ times and smallest AHP peak sizes of all C-fibers (*Table 1*). C5 neurons did not fire AP trains or respond to mechanical stimuli and had the most depolarized RMP (*Table 1*). This group shares characteristics with mouse tyrosine hydroxylase (TH)-positive or vGlut3^+^ neurons (Patil et al., 2018), which are classified as C-LTMR (Usoskin et al., 2015b). C5 neurons could also be cooling units, which have been previously identified in DRG neurons and display fast AHPs and shallow AHP peaks (Djouhri et al., 1998b; Fang et al., 2005).

### A-fiber TG neurons in common marmosets

This section presents the characteristic properties of A-fiber-containing NHP TG neuron subgroups with RMP above -40 mV, which is the main quality control parameter (Djouhri et al., 1998b; Fang et al., 2005).

<u>A1 Neurons</u>: From 32 A1 neurons, 12 were too large for proper compensation. The A1 group exhibited intermediate-duration APs with a “deflection” on the falling phase (*Fig 5A*). Approximately 35% (13/32) of A1 neurons fired AP trains (*Fig 5B, Table 1*). I_A_ currents were recorded from 10 A1 neurons (*Fig 5C*). A1 neurons were the only group of A-fibers that did not respond to mechanical stimulation, a key difference from the A2 group, which showed MA currents (*Fig 5D*). Otherwise, A1 and A2 neurons had nearly identical AP characteristics.

<u>A2 Neurons</u>: Similarly to A1 neurons, the A2 group included the largest neurons, most of which (18/21) had capacitances greater than 100 pF, making them difficult to properly compensate on the Axopatch 200B system. Among the A-fiber groups, A2 neurons had the highest activation thresholds for MA currents and produced the smallest amplitude currents (*Figs 2A, 5D*), with no statistically significant differences in MA characteristics from the C4 group of C-fibers (1-way ANOVA; F (4, 66) = 9.596; n=12 and 21; *Table 2, Figs 2A, 2B, 2C*). I_A_ currents were recorded from 10 of 21 A2 neurons (*Fig 5E*). Given the properties of A1 and A2 neurons, both groups likely represent two types of A-high-threshold mechanoreceptors (A-HTMR) (Patil et al., 2018).

<u>A3 Neurons</u>: The A3 group was characterized by fast-duration APs that lacked the “deflection” seen in C-fiber neurons and A1 and A2 groups (*Fig 6A*). They had a distinct AHP phase without a notable AP undershoot, and their AHP and AHP peak were the shortest among all 10 groups (*Table 1*). Apart from one neuron that fired two APs, A3 neurons did not fire AP trains (*Table 1, Fig 6B*). Of the fast-duration AP groups (A3-A5), A3 neurons exhibited the smallest amplitude MA currents and the fastest decay kinetics (*Table 2, Fig 6C*). I_A_ currents were recorded from 5/9 A3 neurons (Fig. 6D). The properties of A3 neurons are similar to those of TrkB^+^ mouse Aδ-low-threshold mechanoreceptor (Aδ-LTMR) neurons (Patil et al., 2018; Usoskin et al., 2015b).

<u>A4 Neurons</u>: A4 neurons exhibited fast-duration APs with slower AHPs (*Fig 7A*). Only one A4 neuron fired two APs, while the others did not fire AP trains (*Table 1*). A4 neurons had the largest MA current amplitudes and the slowest decay kinetics of all MA-expressing neurons (*Table 2, Fig. 2C*).

**Figure 7.**
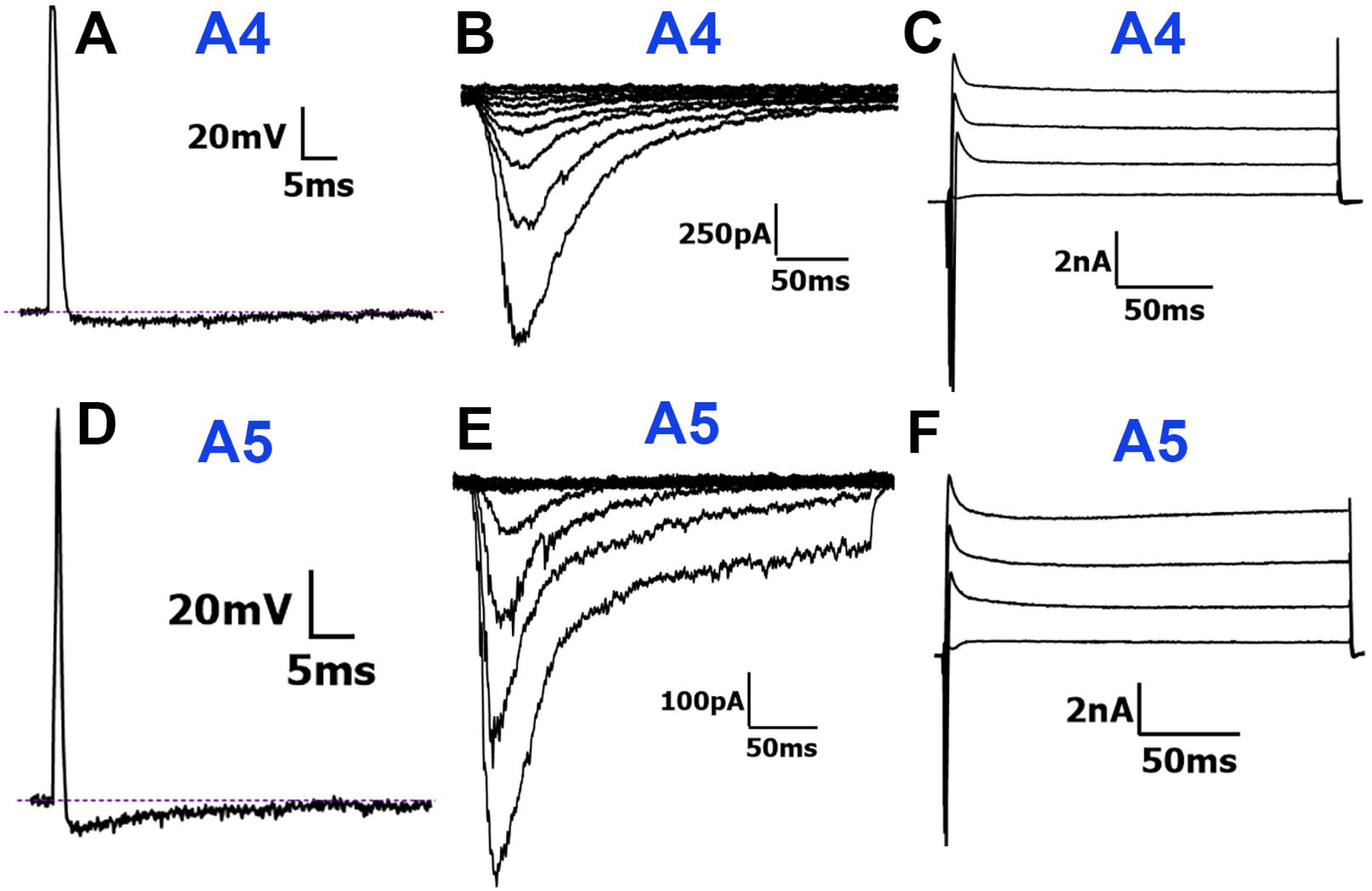
Traces representing current signatures from A4 and A5 NPH TG neuronal groups. **(A)** Representative AP trace belonging to the A4 group. B. Representative MA currents belonging to the A4 group. **(C)** Representative I_A_ belonging to the A4 group. **(D)** Representative AP trace belonging to the A5 group. **(E)** Representative MA currents belonging to the A5 group . (F) Representative I_A_ belonging to the A5 group. Neuronal groups are specified above traces. Scale bars are presented for each panel.

<u>A5 Neurons</u>: A5 neurons also exhibited fast-duration APs but had faster AHPs and notable AHP peaks (*Fig 7D, Table 1*). Apart from two neurons that fired 4 and 7 APs respectively, A5 neurons did not exhibit AP trains (*Table 1*). The MA currents recorded from A5 neurons were similar to those recorded from A3 neurons (1-Way ANOVA; F (4, 66) = 9.596; n=8 and 16; *Table 2, Fig 2*). MA currents in A4 and A5 neurons had the lowest activation thresholds and higher peak current amplitudes, compared to C4 and A2 neurons, which exhibited high activation thresholds and smaller peak current amplitudes (1-way ANOVA; F (4, 66) = 9.596; P<0.0001 for MA current size and F (4, 66) = 5.240; P=0.001 for MA current threshold; n=8-21; *Table 2, Figs 2A, 2B*). In summary, the A4 and A5 neuron subgroups, with their fast APs, large MA currents with low activation thresholds, align with the properties of Aβ-LTMRs reported in the literature (Patil et al., 2018; Petruska et al., 2000a; Zheng et al., 2019).

### Expressions of sensory neuronal markers in TG of common marmosets

According to electrophysiology data, the proportions of the different NHP TG subgroups are as follows: non-peptidergic C-HTMR (C1) is ∼4%; peptidergic C-HTMRs (C2-C4) are ∼33%; C-LTMR and/or cold-nociceptors (C5) is ∼8%; A-HTMRs (A1 and A2) are ∼29%; and A-LTMR (A3-A5) are ∼21% (*Table 1*). According to single-cell RNA-seq data and previous electrophysiological evaluation of reporter mouse lines, C2-C4, A1 and A2 are CGRP^+^ peptidergic neurons, and a subset of C1-C4 are trpV1^+^ neurons (Patil et al., 2018; Usoskin et al., 2015b; Yang et al., 2022). Accordingly, we further validated the identification of neuronal clusters by IHC through the assessment of known sensory neuronal clusters by defined markers.

Peptidergic TG neurons were identified by CGRP (*Fig 8A*) and C2-C4 neurons with trpV1 markers (*Fig 8B*). We found that 58.7 ± 4.5% of neurons were peptidergic, while 37 ± 3.4% of neurons were trpV1^+^ (*Fig 8G*). Among these, CGRP^+^/trpV1^-^ neurons, marking A-HTMRs, accounted for 30.3 ± 4.8% of all neurons, while CGRP^-^/trpV1^+^ neurons, representing a portion of non-peptidergic neurons, constituted 8.6 ± 1% (*Fig 8H*). Additionally, a subset of peptidergic C-fiber neurons, that were CGRP^+^/trpV1^+^, comprised 28.4 ± 3.2% of the total population (*Fig 8H*). These data indicate that A-HTMRs are about 30% of NHP TG neurons. Peptidergic C-fibers comprise at least 25% of NHP TG.In mouse DRG, non-peptidergic C-fiber neurons are classified into three groups: MrgprD^+^, IL31R^+^, and tyrosine hydroxylase (TH)^+^ (C-LTMR) (Patil et al., 2018; Usoskin et al., 2015b; Yang et al., 2022). Within the MrgprD^+^ group, neurons can be further divided into two subpopulations: trpV1^+^ and trpV1^-^ (Patil et al., 2018). Similarly, trpV1 expression has also been observed in the IL31R^+^ group (Patil et al., 2018). Thus, the CGRP^-^/trpV1^+^ population in NHP TG may include some MrgprD^+^ and IL31R^+^ neurons. Interestingly, we detected MrgprD in 17.3 ± 3.1% of NHP TG neurons (*Figs 8D, 8G*). This shows that C1 group is at least 14% of all NHP TG neurons. Tyrosine hydroxylase (TH), a marker for sympathetic fibers in the facial muscles of NHPs (Hovhannisyan et al., 2023), was found to label only a few neurons in NHP TG (*Figs 8E, 8G*). Considering CGRP^-^/trpV1^+^ expression percentage, it could be predicted that C5 neurons compose at least 8% according to IHC, and they can be combination of C-LTMR and cold-nociceptors.

**Figure 8.**
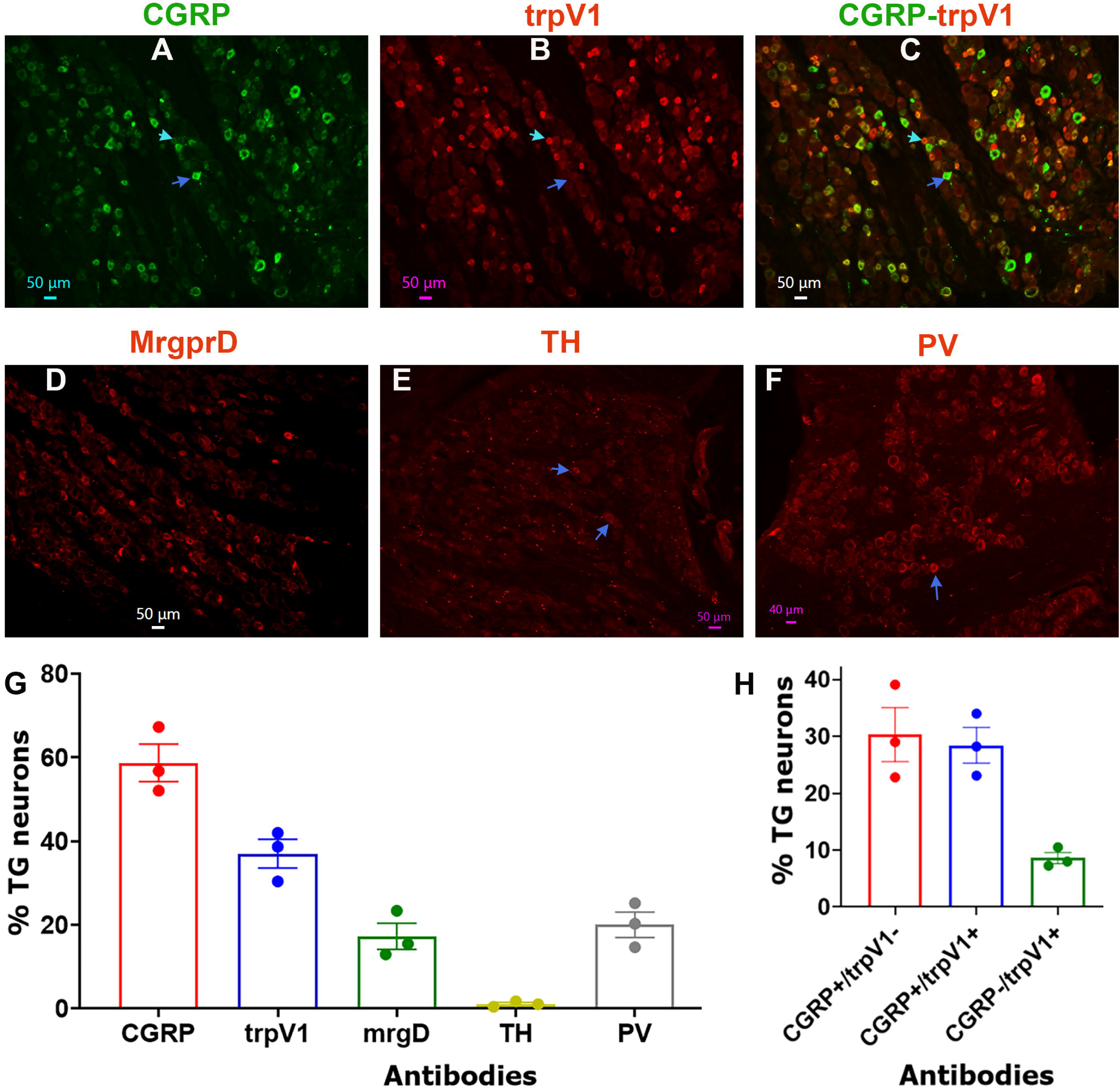
Representation of marker-positive neurons in TG of adult male marmosets. Representative micro-photographs show expression patterns for CGRP **(A)**, trpV1 **(B)**, CGRP+trpV1 **(C)**, MrgprD **(D)**, tyrosine hydroxylase **(E)** and parvalbumin **(F)** in TG of adult male marmosets. Blue arrows on the *panels A-C* indicate CGRP*^+^*/trpV1*^-^* neurons. Cyan arrows on the *panels A-C* show CGRP*^-^*/trpV1*^+^* neurons. Antibodies used and matching colors are indicated. Scales are presented in each microphotograph. **(G)** Bar graphs reflect percentages of marker-positive sensory neurons in TG of adult male marmosets. **(H)** Bar graphs reflect relative percentages of CGRP^+^ (peptidergic) and trpV1^+^ neurons in TG of adult male marmosets. X-axis denotes antibodies for markers. N=3.

A-LTMR neurons are typically identified by markers such as TrkB, TrkC, calbindin (Calb1), and parvalbumin (PV) (Patil et al., 2018; Usoskin et al., 2015b; Yang et al., 2022). Labeling NHP TG sections with TrkB, TrkC, and Calb1 antibodies yielded inconclusive results. However, PV, which is expressed in the majority of A-LTMR neurons (Usoskin et al., 2015b; Yang et al., 2022), was detected in 20.1 ± 3% of NHP TG neurons (*Figs 8F, 8G*). Although PV is also expressed in proprioceptors (Usoskin et al., 2015b), it is important to note that proprioceptive neuron cell bodies in the head and neck are located in the brainstem rather than in the TG. Overall, analysis presented IHC data and presumption of similarity between DRG and TG neuronal markers suggest that ≈25-30% are peptidergic C-HTMR, ≈4-20% are non-peptidergic C-HTMR (including cold-nociceptors, MrgprD^+^ and IL31R^+^) and C-LTMR, ≈30% are A-HTMR, and ≈20% are A-LTMR in NHP TG (*Figs 8, 9*). These conclusions from IHC has similarity with data obtained from electrophysiology result-based clustering.

## Discussion

Isolated sensory neurons have been extensively used to identify functional clusters and investigate pain mechanisms. It is well-established that distinct classes of neurons are responsible for recognizing and transmitting a variety of sensory modalities. Additionally, these neuron groups are believed to have differential roles in the transition to and maintenance of chronic pain conditions. Identifying and characterizing sensory neuronal clusters in NHP TGs enhances the translatability of rodent studies and opens pathways for future research on the involvement of these neuronal classes in both physiological and pathological conditions, particularly chronic orofacial pain. These clusters could also serve as novel therapeutic targets at the whole-cell level (i.e. cell-based therapy).

Using electrophysiological responses to mechanical stimulation, we identified five C-fiber and five A-fiber neuronal groups in NHP TG based on AP characteristics, the presence or absence of AP trains, and transient I_A_ currents (*Table 1*). Additionally, we have used IHC with sensory markers to validate electrophysiology results. *Figure 9* summarizes and schematically represents these NHP TG neuronal groups based on our findings and existing literature. C1 neurons likely correspond to MrgprD^+^ (non-peptidergic) nociceptive neurons, which have been associated with mechanically unresponsive TG neurons in NHP. While MrgprD^+^ DRG neurons in mice are known to express PIEZO2 at higher levels, which is typically found in Aβ-LTMRs. Primate DRGs contain a substantial number of non-peptidergic sensory neurons (Kupari et al., 2021). Interestingly, human DRGs predominantly contain peptidergic neurons (Tavares-Ferreira et al., 2022). C2-C4 neurons share characteristics with traditional peptidergic C-fiber nociceptors (C-HTMRs), which are believed to function as polymodal nociceptors, thermoreceptors and/or silent nociceptors. Since C4 is only neuronal group among C-fiber NHP TG neurons that responded to mechanical stimuli, C4 could represent polymodal nociceptors. C2 and C3 neurons are candidates for thermal or silence nociceptors. C5 neurons resemble those recorded from mouse DRG TH^+^ C-fiber neurons (C-LTMRs), but they may also represent somatostatin-positive (Sst^+^) neurons from the non-peptidergic NP3 group, which is characterized by long-duration APs in (Patil et al., 2018; Usoskin et al., 2015b). Alternatively, C5 neurons could be cooling units, which have been previously identified in DRG neurons and display fast AHPs and shallow AHP peaks (Djouhri et al., 1998b; Fang et al., 2005). Notably, cooling-sensitive TG neurons, which also display fast AHPs, are rare but could also have similar action potential shape (Parpaite et al., 2021). Cold-sensing sensory neurons have been shown to be mechanically insensitive and are rare (Bron et al., 2014; Djouhri et al., 1998b; Fang et al., 2005; Fernandez-Trillo et al., 2020). C-LTMR, trpM8 and NP3 small groups were identified in primate DRG (Kupari et al., 2021).

**Figure 9.**
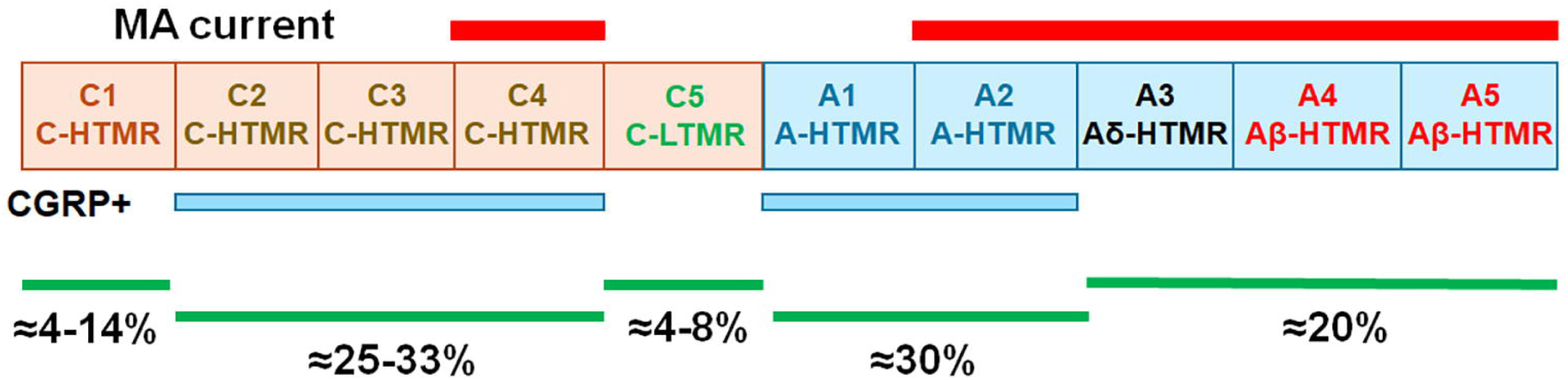
Schematic summarizing results of the study. C-fiber containing NPH TG neuronal groups (C1-C5) represented as the beige boxes. A-fiber containing NPH TG neuronal groups (A1-A5) are the blue boxes. TG neuronal groups responding to piezo-actuating mechanical simulations and exhibiting MA currents are outlined by the red bars and labeled *MA current* above each TG neuronal group. Putative CGRP^+^ groups are outlined by blue lines and labeled *CGRP+*. Relative representations of neuronal groups are based on electrophysiology and IHC data, and are indicated.

While research in pain mechanisms has focused heavily on C-fibers, the functional role of myelinated A-fibers, particularly A-nociceptors, in physiological and pathophysiological pain conditions remains poorly understood. A1 neurons in our study resemble Aδ-nociceptor-like DRG neurons, while A2 neurons also share characteristics with Aδ-HTMRs (Patil et al., 2018; Zeisel et al., 2018; Zheng et al., 2019). Our IHC data suggests that the NHP TG is dominated by CGRP^+^/trpV1^-^ neurons (*Fig 8*). Recordings also indicated a substantial presence of A1 and A2 groups, despite difficulties to maintenance in culture due to their large size. Thus, neurons with deflection of the falling phase of APs, which is hallmark of Aδ-HTMRs (Patil et al., 2018; Zheng et al., 2019), displayed high capacitance, with many neurons exceeding 100 pF. Single-cell RNA sequencing from NHP DRG has identified small proportions of neurons resembling PEP2 and PEP3, which may align with our A1 and A2 findings (Kupari et al., 2021). A3 neurons likely represent Aδ-LTMRs (Patil et al., 2018), while A4 and A5 neurons exhibit features consistent with Aβ-LTMRs (Patil et al., 2018; Zheng et al., 2019). Due to the absence of conduction velocity (CV) measurements in this study, we could not strictly differentiate between Aβ and Aδ fibers. Moreover, Aβ fibers in TG appear more diverse than those in DRG (Harper and Lawson, 1985; Okutsu et al., 2021; Waddell and Lawson, 1990). The relatively small population of A-LTMR neurons identified in NHP DRG through single-cell sequencing aligns with our findings (Kupari et al., 2021).

Strikingly, no fast MA currents were recorded from naïve NHP TG neurons. While proprioceptors and A-LTMRs typically express PIEZO2 and sustain fast, rapidly adapting (RA) MA currents in DRG neurons (Coste et al., 2010; Woo et al., 2015), the absence of proprioceptor cell bodies in the TG could contribute the nonexistence of RA MA currents in NHP TG (Jerge, 1963; Lazarov, 2007). The complete absence of such currents, however, was unexpected. Notably, MA currents recorded from duck TG neurons resemble those observed in NHP TG, with the primary difference being that duck currents tend to have larger amplitudes and slower inactivation kinetics (Schneider et al., 2017; Schneider et al., 2014). The slower inactivation kinetics in NHP neurons may suggest a mechano-nociceptive function (Hao and Delmas, 2010), where slower kinetics allow for greater AP firing during mechanical stimulation (Hao and Delmas, 2010). Additionally, some studies show no difference in mechanical response threshold while others have observed higher thresholds for mechano-nociceptors (Lewin and Moshourab, 2004; Viatchenko-Karpinski and Gu, 2016).

This study employed only one stimulation protocol, though evidence suggests that responses vary depending on the type of mechanical stimulation used, such as stretch or vibration (Poole et al., 2014; Rugiero et al., 2010; Schaefer et al., 2023). Parameters like probe velocity, angle, and diameter, as well as tonicity of solutions, have all also been shown to influence MA current characteristics (Jia et al., 2016; Rugiero et al., 2010; Verkest et al., 2022; Zeitzschel and Lechner, 2024). Patch experiments conducted under substantially uneven resting tension may also alter the kinetics of many channels (Suchyna et al., 2009). Pressure applied by the patch pipette to the membrane to form a giga seal and the size of the patch pipette could affect a cell’s mechano-sensitivity (Cho et al., 2002). In addition, the NPH TG neurons used in this study were cultured in the absence of nerve growth factor (NGF), which is well-known sensitizer of MA currents in sensory neurons (Zeitzschel and Lechner, 2024).

There is a general consensus that the protocol used in this study activates Piezo channels, which account for the MA currents detected in NHP TG neurons (Coste et al., 2010; Parpaite et al., 2021). However, Piezo2 expression in sensory neurons does not perfectly correlate with MA currents (Fernandez-Trillo et al., 2020), suggesting that the molecular mechanisms behind mechanical sensory modalities remain largely unknown. Whether this diversity arises from interactions with non-neuronal cells, the extracellular matrix, or neuron-specific Piezo2 complexes requires further investigation (Sekiguchi and Yamada, 2018). Future experiments, such as back-labeling from peripheral tissues, *in vivo* recordings, and patch-seq, will provide more comprehensive insights into TG neuron types (Parpaite et al., 2021). These advanced methods will improve cell-type identification by integrating information across experimental platforms, ultimately advancing our understanding of sensory neuron function in both health and disease (Lipovsek et al., 2021).

Isolated sensory neurons have been used extensively to identify functional clusters and to study pain mechanisms. It is accepted that distinct classes of neurons are responsible for recognizing and transmission of a variety of sensory modalities. It is also presumed that these different groups of neurons may have differential contributions in the progression of pain chronicity and persistency in many pain conditions (Kupari et al., 2021). Identifications and characterizations of NHP TG sensory neuronal clusters increase translatability of rodent studies; enable future studies on determining involvements of these neuronal classes in physiology and pathophysiology of orofacial chronic pain conditions and could be targeted on whole-cell levels by novel therapeutics.

## Acknowledgments

We would like to thank Drs. Cheryl Stucky (Medical College of Wisconsin; Milwaukee, Wisconsin Wisconsin), Bertrande Coste (Aix-Marseille University; Marseille, France), and Jason Pugh (UTHSCSA; San Antonio, Texas) for their guidance in setting up mechanoclamp. We would also like to thank Dr. Adrienne Dubin from Dr. Ardem Patapoutian’s lab (Scripps Research Institute; La Jolla, California) for providing advice on analyzing mechanically activated currents.

This research work was supported by HEAL Initiative NIDCR/NIH DE029187 (to S.R and A.N.A.); by the National Institute Of Arthritis And Musculoskeletal And Skin Diseases of the National Institutes of Health (NIH/NIAMS) through the NIH HEAL Initiative (https://heal.nih.gov/) The Restoring Joint Health and Function to Reduce Pain (RE-JOIN) Consortium UC2 AR082195 (to A.N.A.); and by A Ruth L. Kirschstein National Research Service Award Individual Predoctoral Fellowship to Promote Diversity in Health-Related Research F31 DE032599 awarded to K.A.L.

## Author Contributions

Conceptualization, K.A.L., S.R., and A.N.A.; Methodology, K.A.L., A.H., and J.M.; Formal Analysis, K.A.L. and A.N.A.; Investigation, K.A.L.; Writing—Original Draft, K.A.L.; Writing—Review & Editing, K.A.L., S.R. and A.N.A.

## Data availability statement

The data that support the findings of this study are available from the corresponding author (K.A.L.) upon reasonable request.

